# Naturalness of forest composition affects vulnerability to climate change and disturbances in Alpine mountain landscapes

**DOI:** 10.64898/2026.01.29.702232

**Authors:** Sebastian Marzini, Katharina Albrich, Alice Crespi, Erich Tasser, Camilla Wellstein, Marco Mina

## Abstract

European mountain forests have been strongly shaped by past human activities, which have influenced their structure and composition. Assessing the natural tree-species composition of current forest landscapes is essential for evaluating their biodiversity potential and for informing management prioritization. High levels of compositional naturalness are often associated with greater ecosystem functioning, but it remains unclear whether forest landscapes that are closer to their potential forest composition are also less vulnerable to future climate change and natural disturbances.

Using a process-based forest landscape model, we quantified the naturalness and the vulnerability to disturbances across a large forested area in the Italian Alps. We developed a spatially-explicit index to evaluate how closely current tree species composition matches potential forest composition. We then simulated future forest dynamics under multiple climate change and disturbance scenarios, using two different initial vegetation conditions on the same landscape – potential vs. current forest – and compared their vulnerability based on changes in species dominance, vegetation structure, and height heterogeneity.

Results indicate that current forests exhibit generally low naturalness compared with their potential forest composition, reflecting historical management and agro-silvopastoral practices. The naturalness score changed depending on elevation across the landscape: forests at low (<1500 m a.s.l.) and high (>2100 m a.s.l.) elevations had low naturalness, while those in the mid-elevation range (1500–2100 m) exhibited medium to high levels of naturalness. Vulnerability to disturbances under climate change differed markedly between the two initial vegetation conditions. Current forest was more susceptible to bark beetle outbreaks, driven by past promotion of Norway spruce and further amplified by warming. In contrast, the potential forest was more vulnerable to wind disturbance, likely due to old-growth characteristics, such as greater height heterogeneity and canopy roughness, that increase blowdown susceptibility.

This study provides the first assessment of forest naturalness using spatially explicit dynamic landscape modelling. Given the projected intensification of natural disturbances under future climates, our findings suggest that promoting more natural forest conditions alone may not guarantee higher resilience to climate-induced disturbances. Instead, management approaches should aim at increasing landscape-level structural and compositional heterogeneity in a balanced manner to minimizing future disturbance vulnerability.

## Introduction

The future of mountain forests is a significant concern today, given their role as vital ecosystems that supply numerous services critical to human welfare and the proper functioning of the biosphere (MEA, 2005; Schirpke et al., 2019). In the Alpine region, these forests are fundamental for timber production, water regulation, carbon sequestration, protection from gravitational hazards such as rockfalls and avalanches, and the maintenance of habitats that sustain biodiversity (Elkin et al., 2013, Seidl et al., 2019). However, ongoing climate change is expected to significantly alter mountain forests in terms of structure, composition, ecosystem functions and produced services (Mina et al., 2017; Albrich et al., 2020b). This trend is likely to increase, as upland areas are warming faster than lowlands at the local scale (Gobiet and Kotlarski, 2020; Pepin et al., 2022). Forests in the European Alps have recently experienced major natural disturbances, such as windthrows and bark beetle outbreaks, likely intensified by changing environmental conditions (Seidl et al., 2014b; Patacca et al., 2022; Marangon et al., 2024).

While climate change contributes to the widespread increase in natural disturbances, current forest structure and composition are also influenced by a long history of management (Conedera et al., 2017). For instance, in the European Alps, mono-specific Norway spruce forests have often been encouraged because of their economic benefits (Vacek et al., 2021), even though they are typically more vulnerable to natural disturbances (Klopčič et al., 2009). As a result of human activity, forest ecosystems have become less natural (Barbati et al., 2007) and are often more unstable and susceptible to changes in the environment (Hartl-Meier et al., 2014; Hlásny et al., 2017). Although it is well established that closer to nature-systems, characterized by higher species richness and greater structural diversity, can better buffer natural disturbances (Brockerhoff et al., 2017; Dobor et al., 2020b; Mohr et al., 2024), some studies have reported that natural old-growth forests and post-abandonment secondary forests may nevertheless be more susceptible to wind disturbance (Schulze et al., 2009; Mantero et al., 2020). Therefore, to our knowledge, it remains unclear whether natural forests are less susceptible to disturbances than forests whose structure and composition have been shaped by past management.

Previous research has examined the naturalness (sometimes also called hemeroby, but see Winter, 2012) of a site by comparing its potential natural vegetation (PNV) to the currently observed composition through a range of different approaches (Strona et al., 2016; Bončina et al., 2017). PNV is a widely applied yet debated concept in ecological research (Chiarucci et al., 2010), typically defined as the vegetation that would occur naturally in a given area under stable environmental conditions, assuming the absence of anthropogenic influences and significant environmental changes (Somodi et al., 2012; Yaynemsa, 2022). Other authors, however, argued that PNV refers to the hypothetical dominant vegetation currently associated with a specific area incorporating both anthropogenic influences and natural changes, thus clearly differing from the concept of primitive or undisturbed vegetation (Loidi et al., 2010; Somodi et al., 2021). This potential vegetation composition is usually determined by using floristic surveys, vegetation mapping (De Keersmaeker et al., 2013; Lillesø et al., 2024) or by statistical modeling tools (Gutierres et al., 2018; Qu et al., 2024). Although this concept has attracted attention, research on naturalness in forest ecosystems has been limited (Myllymäki et al., 2024), particularly in Alpine regions. A research study conducted in Switzerland showed that 45% of forest within inventory sampling plots are ‘not natural’ and therefore more susceptible to disturbances (Scherrer et al., 2023b). The same authors also showed that these forests are less effective at providing current and future protection against gravitational hazards (Scherrer et al., 2023a).

To our knowledge, there are no studies that have used process-based models to evaluate forest naturalness in Alpine forest landscapes, taking into account both spatial detail and future scenarios of climate and disturbances. Assessing the degree of naturalness in current forests through spatially explicit analysis would provide valuable insights for effective management and conservation strategies. Moreover, while managed forests with simplified structure and composition are recognized as being highly vulnerable to future threats (Seidl et al., 2017), the degree of susceptibility for unmanaged forests – whose area has been increasing in the Alps due to land abandonment and reforestation (Schnitzler, 2014; Garbarino et al., 2020; Anselmetto et al., 2024), and may expand even further under new policies like the EU Nature Restoration Law – remains unclear. This is especially true when comparing them to an idealized natural forest landscape that has developed without human intervention. Therefore, evaluating the vulnerability of natural forests under future conditions, in comparison to present forests that bear the effects of previous management practices, can yield valuable information to inform management strategies. In this context, forest landscape models are useful tools to investigate such dynamics.

Here, we applied a process-based forest landscape model to assess the naturalness of a mountain forest territory in the European Alps and to quantify its future vulnerability to climate-driven disturbances given its current naturalness and management legacies. Using a novel, spatially-explicit approach based on the degree of naturalness of forest composition, we first examined how much current vegetation influenced by past management deviates from the potential natural composition of the area. We then analyzed the susceptibility of the current landscape to both climate change and the main natural disturbances occurring in the Alps (i.e., windstorm, bark beetle), compared to an hypothetical natural landscape, assessing the effect of climatic factors as well as forest composition and structure on disturbance impacts. To do so, we addressed two research questions: (a) What is the current degree of naturalness of Alpine forest landscapes given their past management legacies? (b) How does the future susceptibility of managed and potential natural forests to disturbances differ under climate changes? We hypothesize that: (1) today’s forests differ from those that might naturally exist, displaying a low level of naturalness – primarily because Norway spruce has been extensively planted over recent centuries, regardless of elevation, and that (2) today’s forests left unmanaged are more susceptible to disturbances than natural forests, as they have less capacity to absorb such impacts.

## Materials and methods

### Study area

The study area is located in the Eastern Italian Alps (Figure 1), in the northernmost region of Italy, South Tyrol (E 010° 35’ 51.03“; N 46° 37’ 25.01”). It partly embeds the upper part of the Venosta valley (*Vinschgau* in German) covering a total of 30,426 ha. This pilot area is a complex mountainous region consisting of three lateral landscapes: Val Mazia/Matsch in the north, and Val Solda/Suldental and Val Trafoi/Trafoital in the south, bordering Switzerland to the west and Lombardy to the south. The study area has an elevation range between 850-2650 m a.s.l. and it has a dry-inner Alpine climate (Obojes et al., 2018) with annual precipitation of 729 mm and average annual maximum and minimum temperature of 7.01 °C and -2.08 °C respectively, for the period 1980-2020 (see Climate and data scenario section).

**Figure 1.**
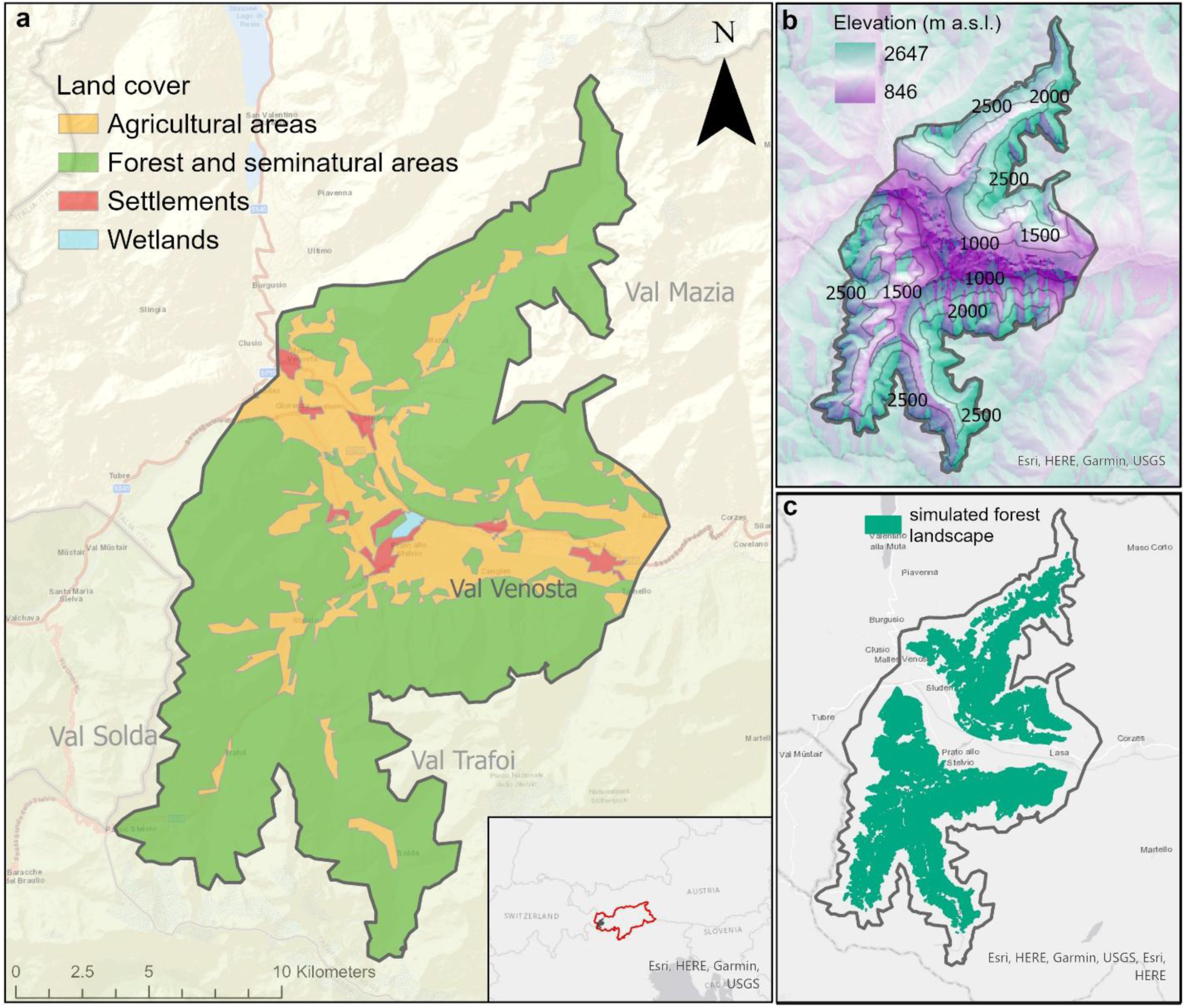
The study area in the upper Venosta valley within the region of South Tyrol: (a) dominant land use/land cover categories, (b) elevation distribution, (c) forested area simulated in this study.

Within this area (hereafter referred as study landscape), 12,589 hectares are covered by forest, predominantly consisting of coniferous species. In the northern part, European larch (*Larix decidua* Mill.), Swiss stone pine (*Pinus cembra* L.; hereafter stone pine), and Norway spruce (*Picea abies* (L) H. Karst.) grow in mixed or pure stands at elevations above 1,600 m a.s.l. In the southern portion, dominant Norway spruce is also accompanied by Silver fir (*Abies alba* Mill.), whose distribution has been limited due to intense browsing pressure from common ungulates such as roe deer (*Capreolus capreolus*) and red deer (*Cervus elaphus*). At lower elevations, Scots pine (*Pinus sylvestris* L.) and Austrian black pine (*Pinus nigra*) – the latter introduced during the last century through reforestation (Aimi et al., 2006) – are present. Near the valley bottom (850-1100 m a.sl.), broadleaf species such as pubescent oak (*Quercus pubescens* Willd.), and black locust (*Robinia pseudoacacia* L.) can also be found. These mountain slopes have been long shaped by human intervention, particularly through grazing and timber production, decreasing overall forests diversity and structural complexity. In recent years, a trend of land abandonment prevalent in the Alpine region has contributed to the expansion of forest area (Kindermann et al. 2023; Tasser et al., 2024; Baglioni et al., 2025). Forest management emphasizes sustainable silviculture, using small gap and strip cuttings on Norway spruce stands, along with selective harvesting and thinning to encourage natural regeneration. The main natural disturbances impacting these forests are windstorms and bark beetle outbreaks, with the incidence of the latter increasing in recent years.

### Simulation model

In this study, we used the forest landscape model iLand, a process-based model that simulates the dynamics of forest ecosystems with a spatially-explicit approach (Seidl et al., 2012; Rammer et al., 2024). Environmental conditions and resource availability (i.e., soil characteristics, nitrogen availability, carbon) together with climate drivers (i.e., daily temperature, precipitation, solar radiation, vapor pressure deficit, CO_2_) are simulated on the entire landscape over a spatial grid of 100×100 m cells (Braziunas et al., 2021; Dobor et al., 2024). Primary production is modelled for individual trees and determined by competition status and available light, which accounts for shading from trees taller than 4 m (Holzer et al., 2024). Tree demographic processes such as growth, regeneration, and mortality are subject to environmental drivers and are partly stochastic. Tree mortality, for instance, is not entirely deterministic but emerges probabilistically, with factors such as maximum age or size, carbon starvation, and disturbances, whereas simulated regeneration encompasses seed production, dispersal, and establishment. iLand also includes sub-models that allow simulating natural disturbances spatially yet using a process-based approach (Seidl et al. 2017). In this study, we focused on wind and spruce bark beetle (*Ips typographus*), which are the main disturbance types across the Alps (Schelhaas et al., 2003; Seidl et al., 2014b). Wind disturbance results from the forest’s structure (e.g., tree stability, exposed edges, and gaps) as well as specific tree species’ resistance to uprooting and stem breakage. These factors influence both the effects and spatial distribution of wind disturbance(Seidl et al., 2014b). Bark beetle disturbance depends on beetle phenology – such as population dynamics, dispersal and colonization – but also host availability and health (Seidl et al., 2017); the intensity of outbreaks is also directly related to simulated climatic conditions. iLand has been extensively applied in Europe for a variety of research studies such as investigating forest disturbances (Thom et al., 2017; Sommerfeld et al., 2021; Dobor et al., 2024; Holzer et al., 2024), forest management (Rammer and Seidl, 2015; Dobor et al., 2020a; Zimová et al., 2020), and functional ecology (Albrich et al., 2020a, 2023; Dollinger et al., 2023). The model has been used at the local, regional and international level to study fire disturbances and post-regeneration dynamics (Braziunas et al., 2018; Hansen et al., 2020; Turner et al., 2022), and forest landscape restoration (Kobayashi et al., 2022; Dollinger et al., 2024).

### Climate data and scenarios

Historical baseline climate was integrated in the model using temperature (minimum/maximum) and precipitation data at daily temporal resolution from 250-m observation-based gridded datasets (1980–2020; Crespi et al., 2021; Mina et al., 2025), resampled to 100-m using a bilinear interpolation. Daily solar radiation was derived from the daily downward surface shortwave flux (DSSF) product available on the LSA-SAF system (https://landsaf.ipma.pt) and downscaled from the original 3-km grid to 250-m resolution by a regression kriging based on elevation, slope steepness and its orientation as described in Castelli et al. (2023). Since DSSF product started in 2004, a temporal resampling was applied to extend the daily time series back to 1980 (Leidinger et al., 2018). Vapor pressure deficit (VPD) was calculated from daily temperature and relative humidity, the latter obtained from the E-OBS dataset (0.1° resolution; Cornes et al., 2018), and statistically downscaled using the *KrigR* package and elevation as covariate (Kusch and Davy, 2022). Mina et al. (2025) provides a detailed account of how baseline climate was parameterized in the study landscape.

Future climate scenarios were retrieved from the daily Euro-CORDEX projections until 2100 (Jacob et al., 2014) based on standard Representative Concentration Pathway (RCP) emission scenarios. We selected six combinations of global and regional climate models from the whole ensemble available which best represent the projected range of long-term (2071-2100) seasonal changes, as spatial averages over the Venosta valley, in temperature and precipitation with respect to historical baseline 1980-2020 (Table 1, Figure 2, Appendix S1: Table S1, Appendix S1: Figure S1). Specifically, we selected three RCP 4.5 and three RCP 8.5 scenarios, which we renamed from CC1 to CC6 according to projected increases in summer temperature at regional scale. These scenarios demonstrate both moderate and strong seasonal differences in rising temperatures and changing precipitation patterns, visible within each RCP as well as across different RCPs (Appendix S1: Figure S2). Further details about climate scenario preparation are provided in Appendix S1.

**Figure 2.**
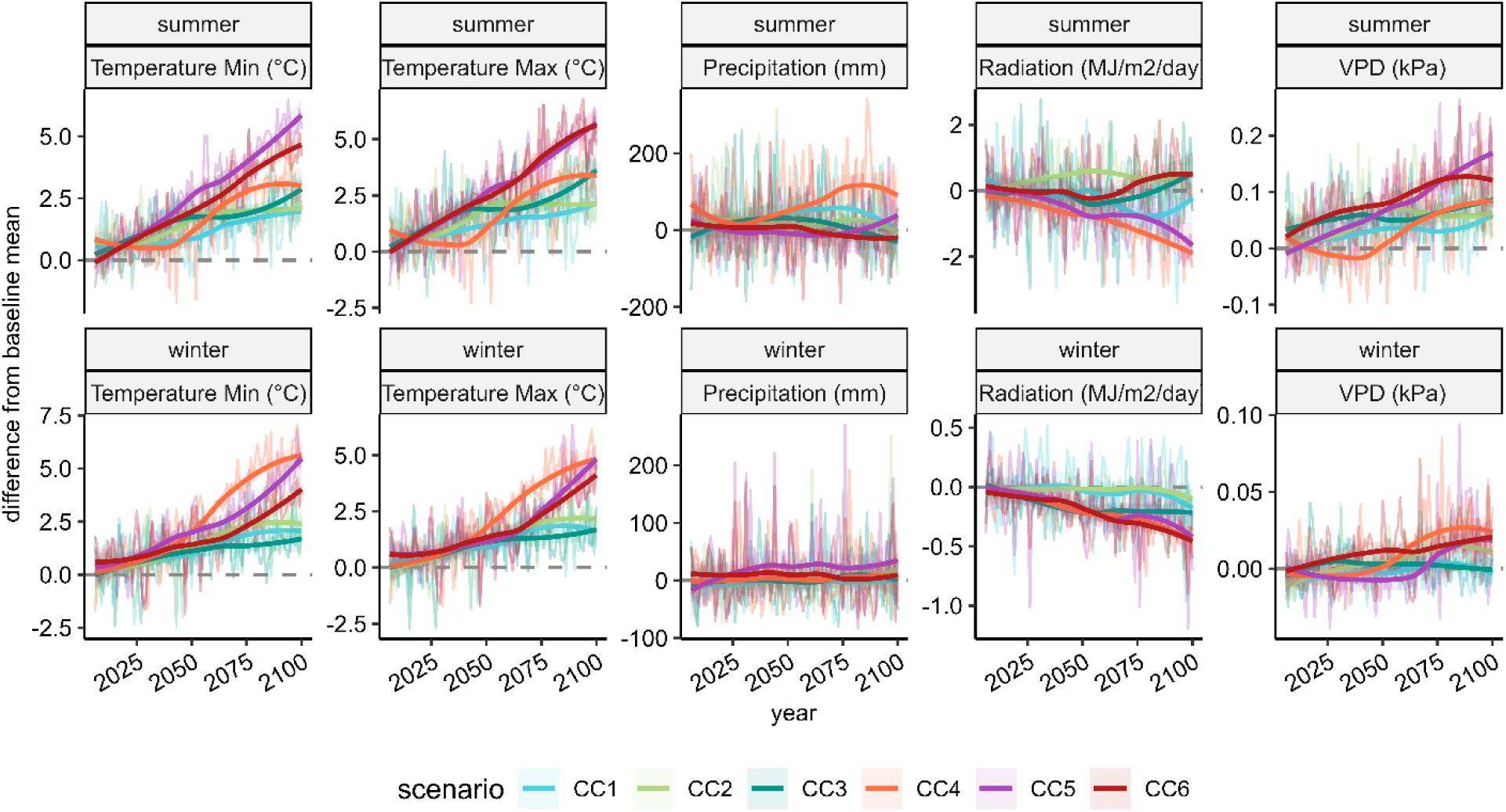
Future changes in the climate scenarios for the Venosta landscape. The figure shows differences to historical baseline climate for summer (J-J-A) and winter (D-J-F) regarding temperature, precipitation, vapor pressure deficit, and solar radiation. Trend lines were smoothed by a loess function (bold). Horizontal black lines indicate no differences to baseline.

**Table 1.** Mean differences of summer (S) and winter (W) temperature and precipitation between the period 2071–2100 and baseline (1980–2020) for the six climate change scenarios used in this study.

| Scenario | RCP | Temperature change (C°) | Precipitation change (%) |
| --- | --- | --- | --- |
| CC1 | 4.5 | S +1.95 / W +1.74 | S -4.91 / W +14.32 |
| CC2 | 4.5 | S +2.29 / W +1.94 | S +8.39 / W +10.40 |
| CC3 | 4.5 | S +1.44 / W +2.46 | S +6.11 / W -4.92 |
| CC4 | 8.5 | S +4.69 / W +3.10 | S +10.08 / W +44.28 |
| CC5 | 8.5 | S +3.78 / W +4.67 | S +27.69 / W +2.24 |
| CC6 | 8.5 | S +3.08 / W +4.66 | S +5.05 / W -8.35 |

To simulate wind disturbance, we retrieved data from the Integrated Nowcasting through Comprehensive Analysis (INCA) system of GeoSphere Austria (GeoSphere Austria, 2021), developed to cover mountain terrains over the eastern Alps (Haiden et al., 2011). INCA provides wind hourly gridded data (wind speed and wind direction) with a spatial resolution of 1 x 1 km. Following Seidl et al. (2014a), we used these data to create a topex-to-distance map to scale wind effect over the landscape in relation to wind exposure given the local topography. To reproduce wind events during the simulations, we generated a single series of wind data, common to all climate scenarios, by randomly resampling monthly maximum wind speeds from the INCA dataset (2012-2020; Mina et al., 2025), and the resampled values were applied to cover the entire simulation period (2023-2200). We opted for resampled INCA data instead of Euro-CORDEX products due to the limited availability of hourly wind simulations in the latter and its coarser spatial resolution, which tends to underrepresent localized conditions, especially for short-duration high wind speeds.

### Model initialization

Our study landscape was set up following the procedures outlined in Mina et al. (2025) , which thoroughly describes all steps involved in initializing, calibrating, and evaluating iLand within the Venosta landscape. Soil characteristics such as depth and texture, as well as available nitrogen were derived from regional forest type maps (Autonomous Province of Bolzano/Bozen, 2010). Initial carbon pools such as soil, coarse woody debris and litter carbon were obtained from the Italian National Forest Inventory (Gasparini et al., 2022), which were spatially imputed to each 1-ha grid cell according to forest type, elevation, slope, and aspect. Forest stand data were collected from local forest management plans that included information such as standing volume, stand age and species shares; these were used to initialize the current forest (12,589 ha) running a spin-up simulation. In iLand, this approach is used to recreate the present vegetation state by modelling long-term succession dynamics, taking into account the effects of previous management actions for each stand, until the resulting forest trajectory closely matches the reference values (Thom et al., 2018). The landscape recreated as a result of the spin-up simulation (see Mina et al. 2025 for details about this procedure) was used as a starting point for simulations into the future and it represents our *current forest* (hereafter CF).

To compare the current forest to a hypothetical natural forest, we generated a potential near-natural forest landscape using iLand. While CF represents a forest landscape embedding the legacy of historical management practices (Mina et al., 2025), this *potential forest* (hereafter PF) represents an unmanaged system driven solely by natural conditions. For generating PF, we included in the model the tree species whose natural range occurs in our study landscape according to the species distribution maps provided within the European Forest Genetic Resources Programme (EUFORGEN, Caudullo et al., 2017). This resulted in 19 indigenous species excluding those whose natural spatial domain did not reach our area or that were favored by human activities in our study landscape (Appendix 1: Table S2). We ran a simulation from bare ground for 1500 years under historic climate conditions and including wind and bark beetle disturbances, until forest composition reached a stable state, thereby accounting for succession dynamics among these selected species that can potentially establish and grow in the landscape. Baseline climate and wind were both derived by resampling their historical data series (climate: 1980–2020; wind: INCA 2012–2020), reflecting the assumption of no climatic changes across the simulation horizon. A snapshot of this potential forest was then obtained by averaging the results from the last 100 years of this long-term simulation.

### Simulation experiment and analysis

We used the two different starting forest conditions (CF and PF) to compute a naturalness score that allows us to evaluate forest compositional naturalness of the present forest in a spatially-explicit manner across the landscape. To calculate the naturalness score, we used the PF as a reference for the hypothetical natural composition of our forest landscape. For each iLand grid cell (1 ha) within the two different forest conditions, we classified tree species into four categories in relation to their simulated presence (species share in terms of basal area) within the cell: 1) Dominant for values between 40-100%; 2) Supplementary for values between 10-40%; 3) Accessory species for values between 1-10%; 4) Not-present for share <1% (i.e., nearly absent). Subsequently, for each pixel in the landscape, we calculated the naturalness score (*NAT*_*score*_) following the approach of Scherrer et al. (2023b). The score reflects the difference in the amount of tree species present in both the PF and CF landscape conditions, weighted according to the dominance category of each species, as defined in the following equation:

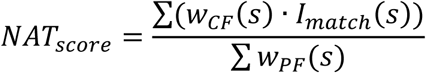

where *w*_*CF*_(*s*) and *w*_*PF*_(*s*) are the weights assigned to CF (observed) and PF (expected) species *s* in relation to the dominancy category (1 if dominant, 0.5 if supplementary, 0.1 if accessory, and 0 if absent), while *I*_*match*_(*s*) was a boolean returning i) 1 when the observed species belongs to the same dominance category of the expected one, or ii) 0 when no match is found. The *NAT*_*score*_was computed by dividing the sum of the weights of the observed species that shared the same dominance categories as the expected species, by the total sum of the weights assigned to the expected species based on their dominance category. The resulted values ranged between 0 and 1, with increasing values related to higher correspondence of the CF respect to PF. Differently from other studies that investigated compositional naturalness for individual sites, our calculation of the *NAT*_*score*_at the pixel level allowed us to spatialize it across the landscape (but see section 4.3 for a discussion using a model-generated naturalness instead of field- or expert-based approaches).

Successively, we used the two distinct forest conditions (CF and PF) as a starting point for our simulations into the future to assess their successional dynamics under different climate scenarios and disturbance regimes. We deliberately excluded simulating future forest management interventions to avoid introducing an additional factor that might have skewed our comparison, which was centred on naturalness and natural disturbances. We started our simulations from 2023, assuming only minor changes in CF between 2020 forest data and the simulation starting year. We simulated forest dynamics from 2023 until 2200 under seven climate scenarios (one continuation of baseline climate and six climate change) including natural disturbances. For each scenario, we run 10 replicates to account for stochasticity, resulting in a total of 140 simulations (2 landscape versions x 7 climate scenarios × 10 replicates). Future climate series for the continuation of baseline climate were generated by resampling (with replacement) of climate data from the period 1980-2020 thereby reflecting present climatic conditions as well. As climate projections were available until 2100, we generated the 2101–2200 series by resampling values from the period 2070–2100. assuming a stabilization of the climatic condition by the end of the century. Wind events were implemented using the time event feature in iLand (Dobor et al., 2024). A single, fixed series of wind events was used for all climate scenarios, with identical timing, wind speed, and wind direction in every simulation. This setup allowed variability in wind-disturbance dynamics to emerge solely from differences in forest stand and climatic conditions, as well as from bark-beetle disturbances affecting stand structure, which interact dynamically with the environment (Seidl and Rammer, 2017). After assessing the variability among replicates (see Appendix S1: Table S2), we grouped and averaged model outputs for each simulation run by scenario.

To evaluate the impact of disturbances, we analyzed yearly wind and bark beetle model outputs aggregated at the landscape level (see Appendix S1: Figure S6 for details of iLand disturbance module outputs), calculating the cumulative extent of disturbed forest at the end of the simulation period and by comparing the results for both PF and CF for the different climate scenarios. To investigate the main drivers of the two natural disturbances considered in this study, we analyzed forest composition (i.e., the proportion of Norway spruce) for bark beetle outbreaks, and forest structure for wind disturbances (Seidl et al., 2023). Differences of wind-related impacts across scenarios were assessed by quantifying vertical structural diversity using the rumple index (RI; Silva Pedro et al., 2017), a metric of canopy structure and surface roughness calculated as the ratio between the canopy surface area and the projected ground surface area (Seidl et al., 2012b; Solano et al., 2022) . Higher RI values suggest increased canopy complexity or lower density due to gaps (Kane et al., 2010); while older stands typically show higher RI, this may not apply in frequently disturbed forests. We further quantified vertical structure heterogeneity by calculating the Shannon diversity Index of the cumulated relative frequency of nine stand height classes of 5-m steps (from 10 to 55 m) for each cell and by comparing the values obtained for PF and CF under the different climate scenarios. Data preparation and analyses were performed using the R Statistical Software (v4.3.0; R Core Team, 2023).

## Results

### Naturalness of the current forest

Our results highlighted clear differences in tree species composition between the idealized natural forest and the current forest (Figure 3). While the set of species present across the landscape was broadly similar, their relative abundances, dominance and distribution along the elevational gradient differed substantially (Appendix S1: Figure S3). The current forest (CF) area was mainly dominated by Norway spruce and European larch, while under near-natural conditions (PF), Scots pine covered a higher portion of the area, especially at mid and low elevations between 600 and 1500 m a.s.l.. Silver fir and Swiss stone pine dominated stands were more prevalent in CF than in PF, where Silver fir was nearly absent.

**Figure 3.**
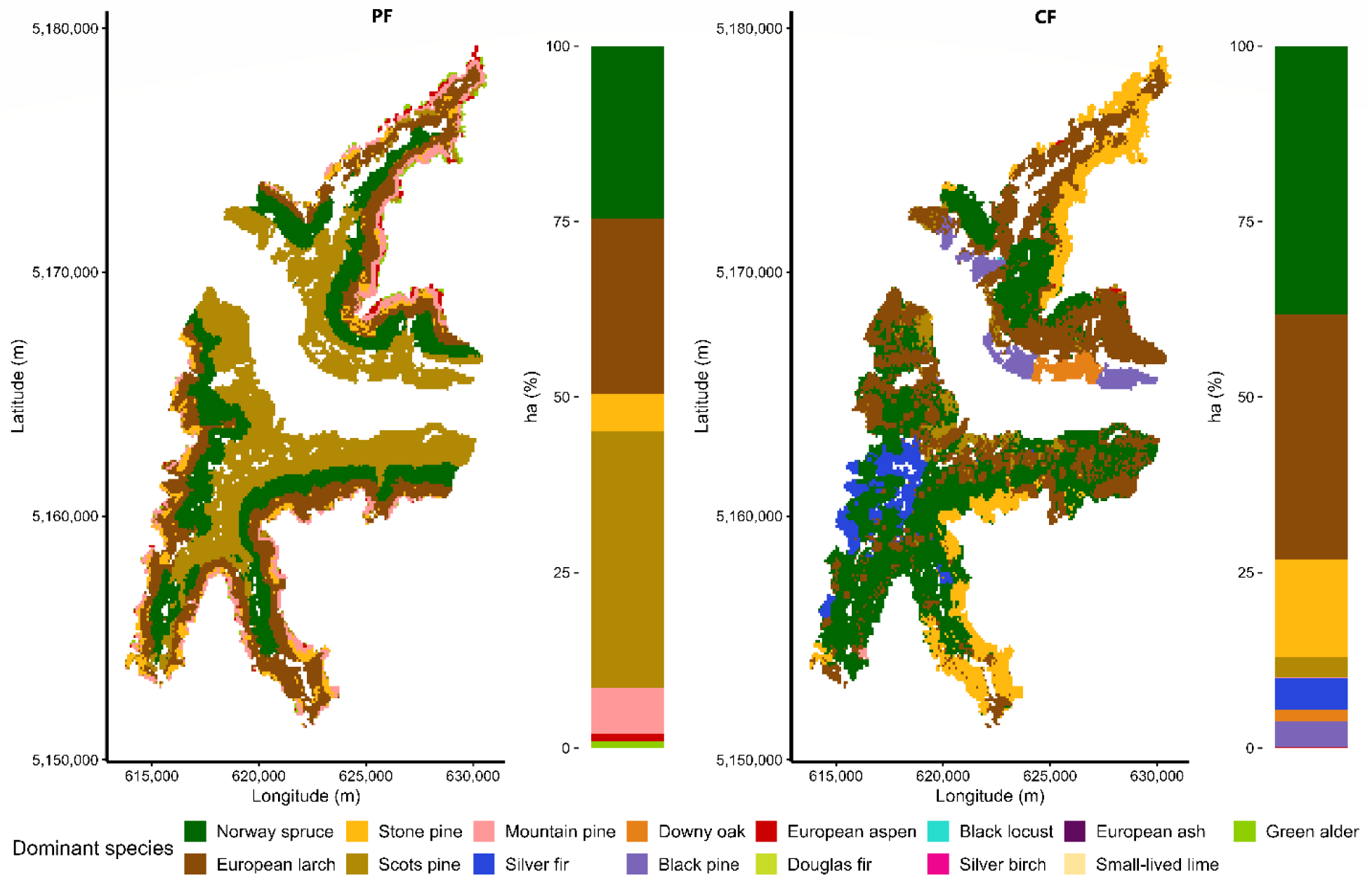
The dominant tree species across the study landscape between the potential natural forest (PF, left) and the current forest (CF, right). Bars on the side of each map summarize the overall dominant species share.

In PF, broadleaved species such as European aspen (*Populus tremula*) and green alder (*Alnus viridis*) – but also montane pine (*Pinus mugo*) – were more abundant at higher elevations (> 1800 m a.s.l.), while in CF the only dominant broadleaved species was pubescent oak (Quercus pubescens Willd.) limited to a small area near the valley bottom. In PF, species distribution exhibited a clear elevation gradient, with species occupying distinct altitudinal zones according to their ecological requirements. The difference was particularly notable for Norway spruce: in PF, this species was found only in upper-montane and subalpine areas (1500–2100 m a.s.l.), while in CF, it occurred throughout the entire range of elevations.

By quantifying the naturalness index at each pixel of the landscape, we produced a *NAT*_*score*_map for the area, reflecting the differences of species dominancy (Figure 4a). The naturalness score map shows that current forests generally have low compositional naturalness, particularly at elevations below 1500 m a.s.l. and above 2100 m a.s.l. near alpine grasslands (Figure 4b). This pattern is mainly due to the presence of species that are completely absent under the PF scenario, such as Black pine, Black locust, Downy oak, European larch, Silver fir, and Douglas fir. Conversely, areas with a higher naturalness were distributed from mid to high elevation ranges, especially on spruce dominated stands between the montane and subalpine belt. By looking at naturalness score categorized into four classes (i.e., high, moderate, low, and very low naturalness) as shown in Appendix S1: Figure S4, we found that all tree species dominating the CF were present across all four classes. However, portions of the landscape with a very low compositional naturalness were mainly dominated by Norway spruce, European larch, Stone pine and Silver fir.

**Figure 4.**
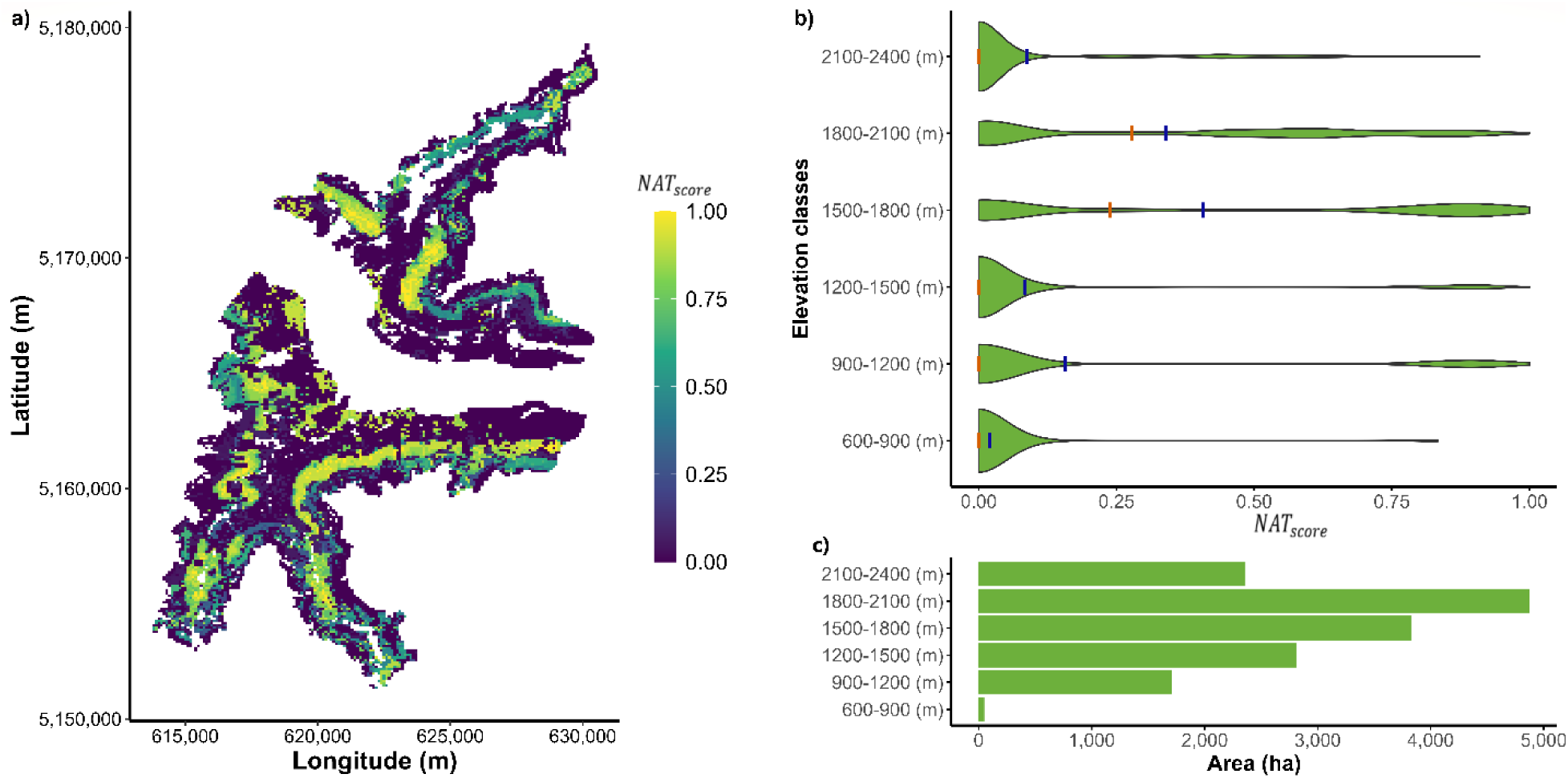
Naturalness score (*NAT*_*score*_) across the study landscape: a) naturalness score mapped spatially in the area, with values going from 0 (very low naturalness) to 1 (high naturalness); b) violin plots showing the distribution of the naturalness score across elevation classes, with mean and median values represented by vertical blue and orange bars; c) horizontal bars showing the extension of the forest area for each elevational class.

### Forest dynamics and susceptibility to climate change and disturbances

Simulations examining forest dynamics under climate change and natural disturbances indicated that climate and disturbance factors had limited impact on species composition at the landscape level. However, our analysis revealed distinct differences in disturbance vulnerability between the two simulation sets initialized with CF and PF (Figure 5). To maintain conciseness, in this section we focused on differences between three selected climate scenarios (baseline, CC1, and CC6), while comprehensive results for all simulated scenarios are provided in Appendix S1 (Figure S5, S6, S7).

**Figure 5.**
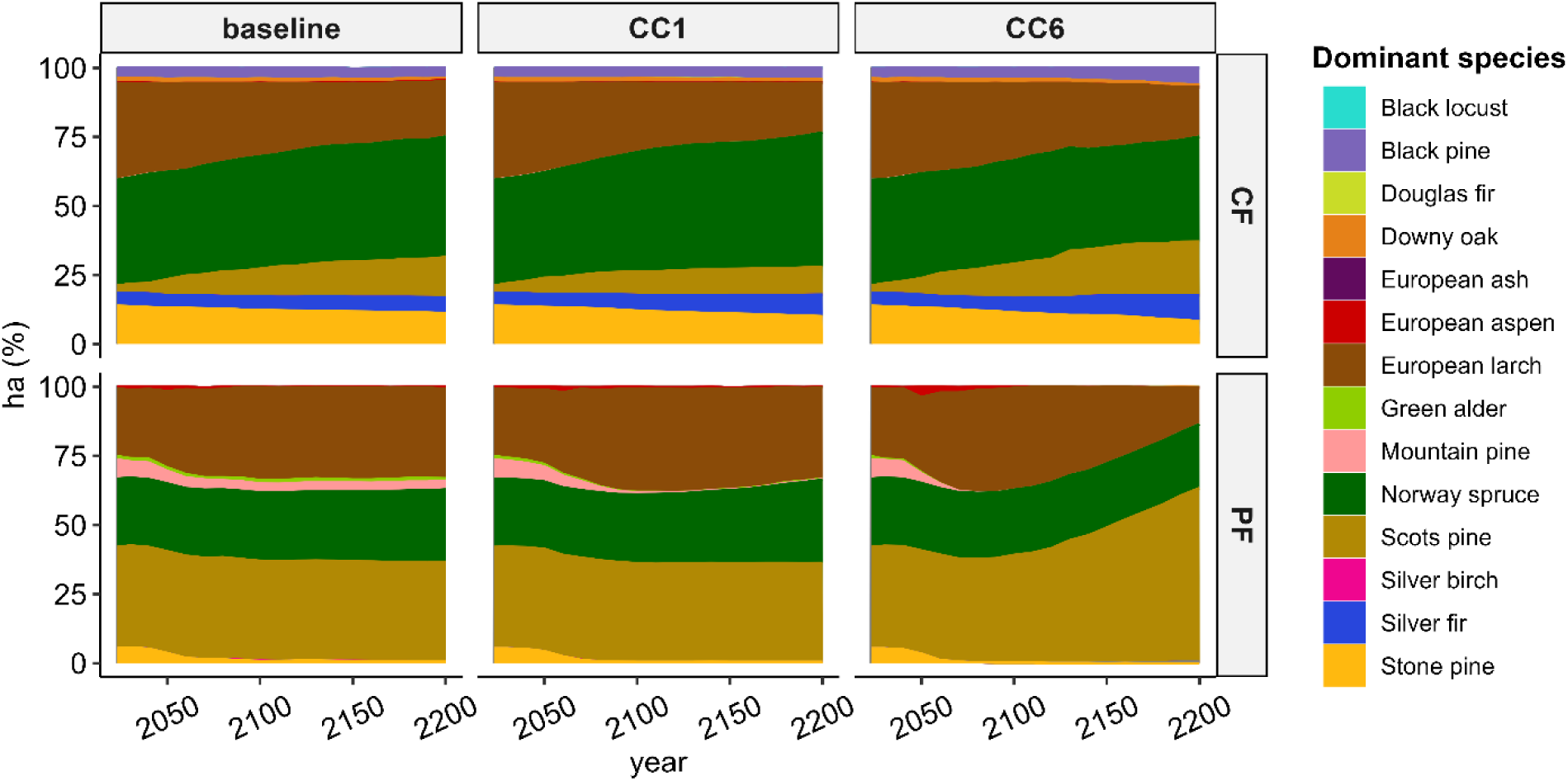
Dominant species share (based on basal area) at the landscape level (in % area) for CF and PF under future climate projections (baseline, CC1 and CC6) until 2200.

In summary, for both initial forest configuration, we did not observe clear trends in species composition changes during the simulation period; however, in some instances, the proportion of dominant species shifted. Generally, conifer species dominated the forest stands, while broadleaves occupied only a small part of the landscape. In PF, Scots pine remained the dominant species in the landscape, expanding more under CC4 and CC5 (Appendix S1: Figure S5) and covering more than the 50% of the entire forest area. Norway spruce and European larch were the second and third most dominant species, with slight shift in their respective trends depending on the climate scenarios. Under the CF initial landscape, Norway spruce remained the dominant species under all climate scenarios, with European larch next, followed by Scots pine, Stone pine, Silver fir, and Black pine. Their distribution varied slightly based on the climate projections.

The impacts of wind and bark beetle disturbances differed between the two initial forest configurations (Figure 6 and Appendix S1: Figure S6). Bark beetle emerged as the most impactful disturbance agent for both initialized landscape and across all climate scenarios. The extent of bark beetle-affected areas was lower in PF compared to CF. Despite this difference, both initialized landscape versions exhibited a clear and marked increase in the proportion of disturbed areas over time across climate scenarios. Specifically, the cumulative damaged area nearly doubled when moving from future projections based on historical climate conditions (14.4% for CF and 8.2% for PF) to the moderate climate change scenario RCP 4.5 (CC1), where the disturbed area rose to 30.7% for CF and 24.7% for PF. This trend continued under the severe RCP 8.5 scenario (CC6), with bark beetle disturbance affecting 67.3% of CF and 60.2% of PF landscape. The strongest impact of bark beetle was however observed under CC5 within the CF initial landscape, where cumulative beetle damage reached as high as 72.4% of the total forested area over the entire simulation period. Similarly to bark beetle dynamics, wind disturbances exhibited a consistent upward trend across future climate scenarios. Under baseline climate projection, wind-related disturbances affected 4.6% of forest under CF and 8.5% under PF. Under RCP 4.5, these values increased to 6.6% in CF and 10.2% in PF (CC1). Under RCP 8.5, the numbers rose even higher, with disturbances affecting 8.7% of CF and 12.2% of PF (CC6). However, in contrast to bark beetle disturbances, the PF version exhibited greater susceptibility to wind events. The most significant wind disturbance impact was found under the CC4 scenario, with approximately 14.3% of the forest area affected by 2200 (Figure S6).

**Figure 6.**
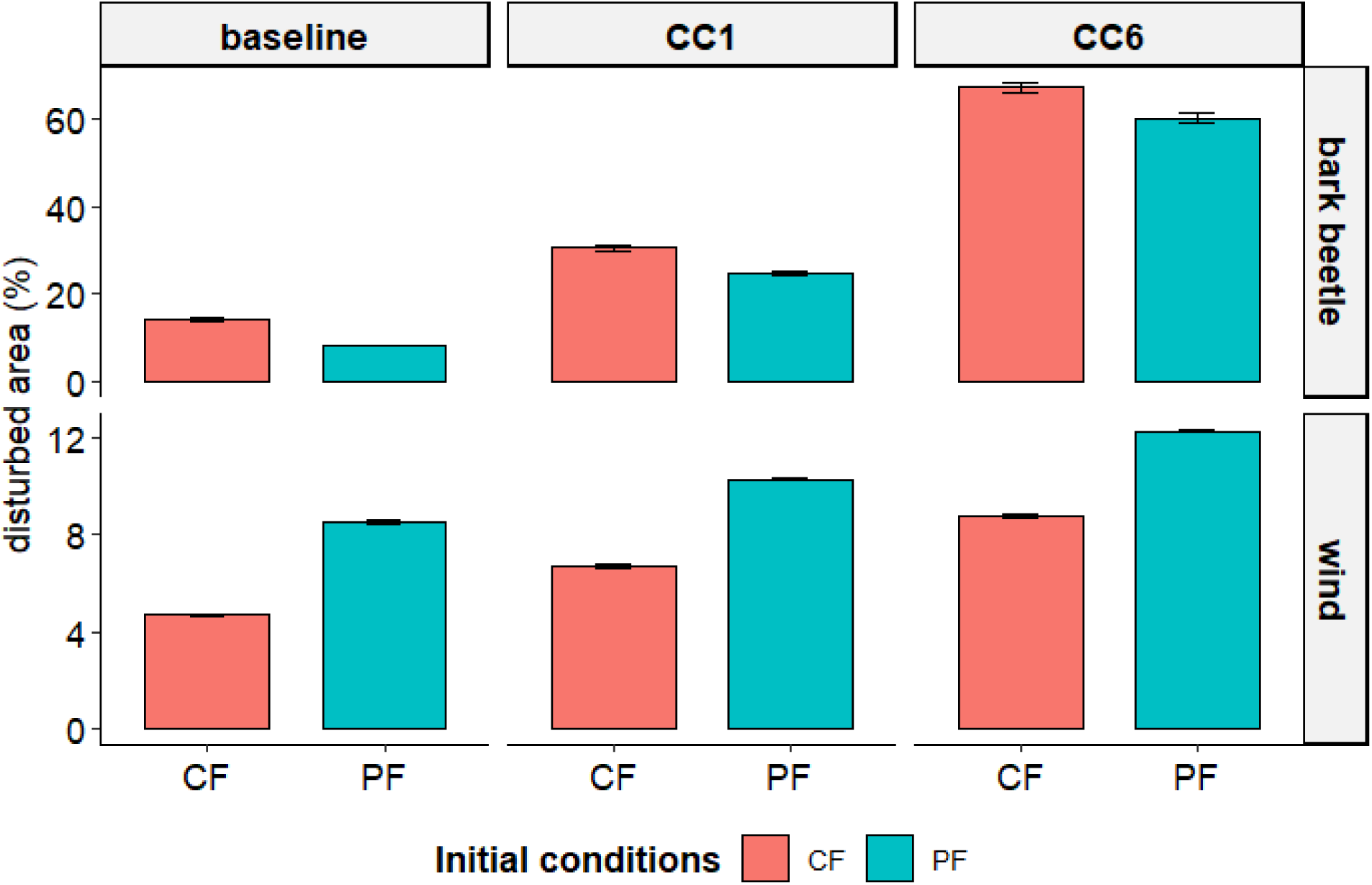
Cumulative damages in terms of percentage of disturbed area for bark beetle (upper panel) and wind (lower panel) on PF (red) and CF (blue) under baseline climate, CC1, and CC6. Error bars show the standard deviation of the replicate mean. Note the different scales of the y axis for the two disturbance agents. Refer to Figure S6 for more details about disturbance outputs.

Forest susceptibility to disturbances is directly affected by forest structure and composition. For bark beetle, the clear main driver is forest composition, specifically the presence of Norway spruce. Generally, bark beetle disturbance was higher in CF than in PF due to the stronger presence of Norway spruce in the forest landscape (Figure 5). As the share of stands dominated by Norway spruce expanded throughout the simulation period under all scenarios, the extent of areas affected by bark beetle disturbance grew accordingly, with this effect being more pronounced under more severe climate change scenarios. Wind disturbance susceptibility is primarily driven by forest structure, in particular the heterogeneity of vertical structure. To investigate this further, we utilized two metrics of vertical structure diversity, the Rumple Index (RI) and the Shannon diversity Index of height classes. The RI showed different distribution between the two initialized forest conditions (Figure 7a and Appendix S1: Figure S7a). PF displayed a highest median of RI values, particularly at the start of the simulation (2023). For this initialized forest condition, differences in RI between the start and the end of the simulation (2200) were minimal, with a slightly higher RI median found only under RCP8.5 (CC6). Instead, under CF the median of the RI was higher at simulation end, particularly under RCP8.5 (CC6).

**Figure 7.**
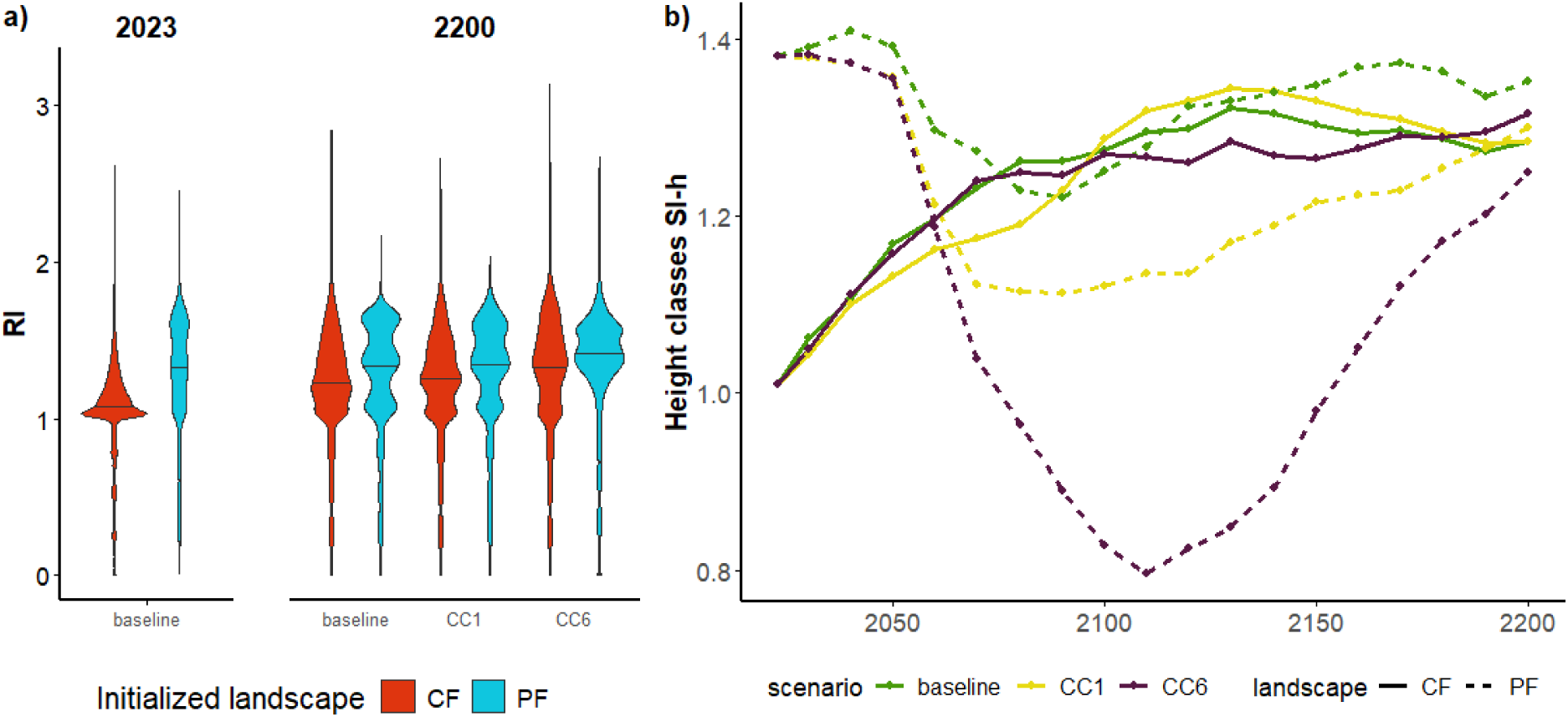
Panel (a) Violin plots showing the rumple index (values of the distribution across the forested landscape extracted at 1 ha resolution) comparing PF and CF at the beginning and the end of the simulation under baseline, CC1 and CC6 scenarios (median: horizontal black line). Panel (b): Temporal trends of the Shannon diversity Index of the height classes (SI-h), representing the evenness of the stand height classes distribution plotted over time with time steps of 10 years. Higher values indicate greater diversity in vertical stand structure as well as the evenness of their distribution.

The Shannon diversity Index of height classes (SI-h) revealed marked differences between the two initialized forest conditions (Figure 7b). At the beginning of the simulation, SI-h was clearly lower in CF compared to PF, indicating that the current forest had a lower vertical stand structure heterogeneity across the landscape compared to a potential forest. During the simulation, SI-h for CF increased steadily, reaching values comparable to PF at the end of the 21st century, and even higher value than PF by the end of the simulation period. This pattern was consistent across all climate scenarios, except under the historical climate, where PF generally maintained a higher SI-h than CF. This trend of increasing SI-h for CF was more pronounced during the first decades of the simulation period and it gradually stabilized over time, except in CC3 where it slightly decreased before the end of the simulation period (Appendix S1: Figure S7). In contrast, SI-h in PF first decreased, then either stabilized or rose slightly by the simulation’s end. Although SI-h trends for CF were consistent across the different climate scenarios – with a general rise leveling off midway through the simulation – PF showed a much steeper decrease of SI-h under the most severe climate projections (CC3, CC5, and CC6). These results suggest that the current forest, when left unmanaged, will developed a higher vertical stand structural heterogeneity across the landscape, potentially raising its vulnerability to wind disturbances, similarly to PF.

## Discussion

Our study presented a novel approach to map the compositional naturalness of forests using dynamic modelling. Our assessment of forest naturalness represents an innovative contribution to the evaluation of naturalness in forest systems by means of spatial-explicit modelling, offering a valuable complement to traditional classification methods through the application of forest landscape models. Our results support our first hypothesis that today’s forests, in our Alpine study landscapes, differ from those that might naturally exist in terms of tree species composition. Despite these general findings, our method highlighted that there are still portions of Alpine landscapes that display high level of compositional naturalness that can be detected by means of spatially-explicit analysis. Our findings also partly support our second hypothesis that current forests, when left unmanaged, are more susceptible to natural disturbances than a potential natural forest. However, this depends on the type of disturbance type considered, as our analysis also revealed that a potential natural forest, characterized by higher vertical structure heterogeneity at landscape level, is more susceptible to windthrow than the current forest.

### Compositional naturalness of Alpine forests

In alignment with findings from research in Swiss forests, reporting a substantial presence of non-natural tree assemblages (Scherrer et al., 2023b), our analysis revealed that a large proportion of our study landscape is currently characterised by low compositional naturalness. This can be mainly explained by a long history of forest management in Alpine areas (Bebi et al., 2017; Conedera et al., 2017), and the fact that many forests that we observe nowadays can be considered semi-natural, therefore resulting from succession dynamics driven by human interventions and expansion due to land abandonment (Tattoni et al., 2017; Agnoletti et al., 2022). This is especially true for Norway spruce stands, the most dominant species in our forest area that was strongly promoted for economic purposes outside their environmental optimum (Johann et al., 2004; Klopčič et al., 2017). By contrast, Silver fir was heavily reduced by cuttings, and today its distribution deviates most from its natural potential (de Jel and Vacik, 2012; Tinner et al., 2013; Coşgun et al., 2025). In our potential forest landscape Silver fir was rarely present as a dominant species, more often acting as a co-dominant across the landscape. Even though the high occurrence and the widespread distribution of Norway spruce observed in the current forest can likely be attributed to the legacy of past management practices, which were embedded in the spin-up simulations to build the initial landscape, the areas dominated by Silver fir may indicate habitats particularly suitable for the species, or represent remnants of its past distribution. Nevertheless, Norway spruce was also present in areas exhibiting greater levels of naturalness, primarily between 1500 and 2100 m a.s.l. across our landscape. This indicates that, despite historical management activities, certain stands within this elevation range continue to exhibit an ideal or near-natural species composition.

For several species, their current occurrence area did not match with what would be expected naturally, indicating a notable divergence from the potential natural species composition. For instance, most of current Norway spruce stands were found on areas potentially dominated by Scots pine, a species that displayed a wider distribution in our simulated potential forest. A larger extension of Scots pine as dominating species would be expected due to its capacity of establishing in drought-prone areas and to its wide environmental plasticity (Rigling et al., 2013; Brichta et al., 2023). This species also acts as a pioneer, capable of colonising disturbed environments and forests where succession is driven exclusively by natural processes. Additionally, the extended dominance of European larch in several portion of the current landscape is likely a result of past land-use practices, as this species has been promoted at low- and mid-elevations to combine timber and grazing functions (Johann, 2007; Schulze et al., 2007; Nagler et al., 2015). Research indicates that larch forests at montane elevations may not be well-suited to present climate conditions, as decreased water availability and increased water atmospheric demand from recent climate changes are affecting larch growth at lower elevations in dry Alpine areas (Obojes et al., 2018). Some of our sites with a low naturalness score were also characterized by the presence of species that were artificially introduced. In our case, black pine plantations at low elevations were established in the last century to increase soil stability and restore mountain slopes after the abandonment of grazing (Vallauri et al., 2002; Mikulová et al., 2019). Currently, these pines are experiencing vitality declines in drought-prone sites (Móricz et al., 2018), and local forest management is working to introduce broadleaved species like native oaks. Lastly, the higher presence of montane pine, green alder, and European aspen at higher elevation in our simulated potential forest is likely related to the absence of management practices that concurred to their strong reduction in the actual high forest areas. While in current managed forests these species have more difficulties to establish and to grow, their distribution is more concentrated around forest stands as shrub encroachments in disturbed and open areas, but their dynamics in the Alps are still under debate (Pisetta et al., 2012; Caviezel et al., 2017).

### Compositional and structural changes affect disturbance vulnerability

Throughout the simulation period into the future, species dominance at landscape level changed only slightly, presenting minor fluctuations among the different climate scenarios. Despite the influence of natural disturbances (discussed further below), the main dominant species remained Norway spruce in CF and Scots pine in PF. Such behavior is common in research involving forest dynamic models because these models are very sensitive to their initialization data. This sensitivity leads to a legacy effect that can influence the future development of forest ecosystems over decades or even centuries (Temperli et al., 2013b; Thom et al., 2018). Compositional changes generally occur more slowly than structural ones and are mainly driven by factors such as nutrient availability, large-scale disturbances, or shifts in climatic niches (Anyomi et al., 2022), none of which were explicitly represented in our modelling experiment. Moreover, the simulated disturbances primarily affect mature trees, thereby preserving species dominance in the younger layers. Consequently, although spruce dominance might be expected to decline more rapidly in CF, its current overrepresentation and sustained recruitment into seed-producing age, even under severe bark beetle disturbance, likely enable its persistence, consistent with Fischer et al. (2015), who found stand composition to remain largely unaffected by bark beetle outbreaks due to intact regeneration layers. Differences in species composition partially explained our results about future disturbance impact. Natural forests are usually seen as systems that are less vulnerable to disturbances (Scherrer et al., 2023b). However, in our modelling experiment we found this to be the case for disturbances agents that are highly specialized (i.e. bark beetle), while this pattern may not hold for more general agents, such as wind, which depends more strongly on stand structural characteristics. These results, however, need to be interpreted within the assumptions and limitations of the modelling framework (see further below). By comparing the simulation set with the two different initial forest conditions, we found a higher vulnerability to bark beetle outbreaks in CF, likely due to the higher share of Norway spruce in the current landscape as a result of past management (Seidl et al., 2018; Sommerfeld et al., 2021; Washaya et al., 2024). In all simulation sets, the infested areas almost doubled under the most severe climate projections (RCP 8.5), and particularly under the CC5 scenario. This climate scenario involves a substantial rise in temperatures – especially during winter – and an increase in VPD. These changes place significant stress on Norway spruce trees and create conditions that allow beetles to thrive, as demonstrated by similar outcomes in studies by Sommerfeld et al. (2021) and Temperli et al. (2013). In the simulation set with the initialized potential natural forest, this effect might be counterbalanced by a higher share of Scots pine and by the distribution of Norway spruce at higher elevations, as forest vulnerability to bark beetle has been found to be lower at high elevation (De Groot et al., 2023).

In addition to bark beetle disturbances, wind-affected areas also increased across all climate scenarios for both of our simulation sets. This trend is likely attributable to reduced tree resistance under severe climatic changes such as those represented by CC3, CC5, and CC6; in iLand, such conditions lead to fewer days with frozen soils that can increase the likelihood of windthrows. This is consistent with the findings of Seidl et al. (2018), who showed that disturbances from wind in an Alpine mountain landscape will likely increase in response to future climatic changes. Interestingly, wind impact was higher in the simulation sets with initialized potential natural forest, no matter the climate we simulated. This might be related to the fact that older forests are particularly prone to windthrow and they display a higher susceptibility due to long-term effects of stand structure and, to a lesser extent, of composition (Harper et al., 2005; Wohlgemuth et al., 2022). Given that the current forest contains more wind-resistant larch and younger spruce stands resulting from previous management practices, it is likely less vulnerable to wind damage than a landscape dominated by mature, late-successional trees, such as in PF. The landscape initialized with these forest conditions exhibited a greater canopy complexity (as indicated by the rumple index), which remained relatively stable throughout the simulation period. In contrast, CF showed a marked increase in canopy complexity during the simulations, particularly under RCP 8.5 climate projections. Old-growth stands like those in PF have generally higher structural canopy complexity (Chamberlain et al., 2021) because of the presence of gaps created by natural successional dynamics (Kane et al., 2010), which in turn can increase susceptibility to windthrow. Additionally, old-growth and post-abandonment forests with high growing stock and advanced stand age are known to be more susceptible to windthrows (Mantero et al., 2020). Older, structurally unstable stands are more likely to collapse during wind events (Schulze et al., 2009; Torresani et al. 2024). While this poses little concern in remote forests with minimal human activity, it is significant in the Alps, where mountain forests provide multiple functions – including protection from gravitational hazards – and are often near settlements that require them to be stable and wind-resistant (Stritih et al., 2024).

The increase of structural stand complexity observed in our simulations with CF could be explained as the forest develops towards with greater diversity of tree sizes and complementarity in crown architectures in the absence of silvicultural interventions, as also shown by Ehbrecht et al. (2021). Intermediate-severity wind disturbances have also been observed to increase stand structure complexity (Peterson, 2019) and this effect might be amplified by climate changes in current forests with a lower complexity due to past management. Conversely, it is interesting that our simulations starting with the potential natural forest showed a decrease of stand-height heterogeneity under all climate change scenarios. This suggests a combined negative effect of climate and disturbances on forest structure, particularly pronounced under scenarios CC3 and CC5, which induced a drastic decline of vertical structure heterogeneity (as indicated by the Shannon diversity Index of height classes) and a delayed in its recovery compared with the other scenarios. However, no clear pattern emerged in relation to the specific climate scenarios or to the different combinations of reduced precipitation, increased VPD, and higher temperatures. Further analyses may therefore be needed to disentangle the climatic drivers underlying these trends.

The initial lower vertical structure heterogeneity and stand canopy complexity observed in CF was likely due to the legacy of past management and the presence of more homogenous and younger stands across the landscape. Silvicultural interventions, such as gap creation or selective thinning, tend to promote vertical stratification by enhancing height differentiation, while also leading to a more homogeneous canopy structure at the stand level, thereby reducing canopy roughness compared to unmanaged forests (Dieler et al., 2017). At the same time, the decreasing stand height heterogeneity in PF due to the presence of fewer but more different height classes, may have contributed to an increased edge effect (Blennow and Sallnäs, 2004), which is strongly associated with wind disturbance impacts. Moreover, a negative effect of high structural diversity on wind susceptibility can result due to differences in species composition (light-demanding vs. shade-tolerant) and height (Wohlgemuth et al., 2022). Additional variables could be considered to better characterize forest structure, such as stem density or stand age, which is often higher in natural forests (Baran et al., 2020). Although we did not analyze stand age directly, the high stand heterogeneity in PF and the observed trends of vertical structure heterogeneity suggest that wind events primarily affected older stands, thereby reducing stand-height heterogeneity. Similarly to the findings of Seidl et al. (2025), our study showed that climate-mediated disturbances are expected to have a strong impact on old-growth systems, but such impacts might be reduced avoiding an unconditional forest aging and by improving stand structure.

### Limitations and recommendations for future research

Our study introduces a novel way to assess mountain forest naturalness using spatial-explicit modelling, but it also has inherent limitations common to dynamic modelling approaches. First, uncertainty remains regarding the accuracy of the simulated potential natural forest without human influence, as no real-world references exist for direct comparison. Our evaluation therefore relied on the model’s ability to reproduce key ecological processes in the resulting landscape, supporting the consistency of the methodology with both the model’s internal framework (Rammer et al. 2024) and the data specific to our study area (Mina et al., 2025). Therefore, the naturalness values presented and discussed should be interpreted as model-based indicators. Second, differently from the approach of Scherrer et al. (2023b), we relied on the analysis of naturalness by using dominant species instead of forest categories or types. We assumed that these variables would better reflect high-resolution changes and help reduce uncertainties arising from further categorizations. However, we did not find a clear reference on which values of species share to use to divide stands into different classes of dominancy. Therefore, to characterize species dominancy, we still used the categories proposed in Scherrer et al. (2023b) but we relied on percentage thresholds that we set arbitrarily by looking at the share of species used to determine the different forest categories in our region. Third, in this simulation experiment we excluded additional variables that could have introduced confounding effects into the interpretation of the results. For example, we did not simulate management interventions into the future, as we aimed at assessing forest successional dynamics resulting and the exclusive impact of natural disturbances in interaction with changing climate. An accumulation of deadwood within the stands due to absence of salvage logging and other silvicultural activities could also play a role in the increase of bark beetle disturbances in the initialized current forest. A comparison of different management regimes in our study area – including alternative management to current strategies – should be addressed in future studies, including specific assessment of the ecosystem services provisioning. Browsing was also excluded as a potential disturbance factor that could negatively impact various forest species, particularly Silver fir (Frei et al., 2024). Moreover, given the influence of climate change on wind disturbances and their interactions with forest ecosystems (Seidl et al., 2017), different wind scenarios might also lead to different results. We intentionally avoided adding this additional layer of uncertainty in the present study, but this aspect could be incorporated in future work.

### Conclusions and implications for management

Our study offers a novel approach for assessing the naturalness of Alpine forest landscapes and it shows that most existing forests differ from natural conditions because of historical management practices. Yet, our analysis showed that forests resembling their potential natural vegetation might not always be the best choice for reducing vulnerability to disturbances caused by climate change. While human activities have increased forest landscapes’ susceptibility to specific disturbance biotic agents like bark beetles, this amplified vulnerability does not extend to more general abiotic agents such as wind, whose impact is more closely linked to the structural characteristics of the forest stands.

In the context of ongoing forest abandonment and natural reforestation in the Alps, leaving current forests unmanaged may exhibit lower resistance with possible negative consequences for ecosystem services provision. Forest management strategies and restoration initiatives in human-influenced landscapes should be meticulously designed at various spatial scales to ensure multifunctionality (e.g., timber, protection, recreation) and support biodiversity (e.g., with a strategic network of non-intervention biodiversity hotspots), while simultaneously enhancing resilience to climate change and the growing frequency of natural disturbances. Despite the considerable complexity involved, studies with forest dynamic models offer valuable insights for evaluating management strategies and envisioning future scenarios for resilient forest landscapes.

## Acknowledgements

SM’s PhD Scholarship was supported by funding from Eurac Research and the Free University of Bolzano/Bozen. MM acknowledge support from H2020-MSCA-IF project “REINFORCE” (grant number: 891671). ST acknowledge funding from the Interreg Italy-Switzerland project “MAP-REZIA” (ID 0200061, CUP D53C24005760007). ET was partly funded by the Austrian Climate Research Program (ACRP, 13th Call) project UNRAVEL (registration number KR20AC0K18081). The authors thank the Forest Planning Office of the Provincial Forest Services of South Tyrol for providing local data to parameterizing the model. The study was co-funded by the open access funding provided by Libera Università di Bolzano within the CRUI-CARE Agreement. Microsoft Copilot was used to improve the language in parts of this manuscript.

## Authoŕs contribution

SM and MM conceived and planned the research, designed the study area, initialized climate and vegetation conditions, and wrote the original draft. SM performed the simulations, formal analysis and visualization. MM provided supervision and funding acquisition. AC collected and processed climate data. KA provided methodological support. ET, CW and KA reviewed and edited the manuscript. All authors contributed to the final version of the manuscript.

## Conflict of Interest Statement

The authors declare no competing interests.

## Data availability

Data that supports the finding of this study (model input files) will be archived in an external repository (e.g., Zenodo) upon the acceptance of the article. A first set of model inputs (biophysical parameters, baseline climate database, digital elevation model, wind data, forest cover area, forest landscape snapshot) for the Venosta study area is already available at https://doi.org/10.5281/zenodo.14886673. The iLand model, including documentation and the full source code, is freely available at https://iland-model.org/ under the GNU General Public License.

## Notes

### Competing Interest Statement

The authors have declared no competing interest.

