## Appendix S1 for "Naturalness of forest composition affects vulnerability to climate change and disturbances in Alpine mountain landscapes"

### Supplementary material for:

***Naturalness of forest composition affects vulnerability to climate change and disturbances in Alpine mountain landscapes***

Sebastian Marzini, Katharina Albrich, Alice Crespi, Erich Tasser, Camilla Wellstein, Marco Mina

### Climate change scenarios

The future scenarios for the required climate variables were retrieved from Euro-CORDEX (Jacob et al., 2014) and post-processed in order to assure the compatibility with the data used for describing baseline climate conditions (Mina et al., 2025). Daily series for 15 temperature and 11 precipitation models for two future emission scenarios (RCP 4.5 and RCP 8.5) were downscaled to 1-km grid by means of observations, covering the period 1970-2100. To optimize the number of scenarios, we compared the projected temperature and precipitation changes in winter and summer over the entire South Tyrol for individual models by emission scenario (Figure S1) to select a number of model/scenario combinations to best covering the range of changes.

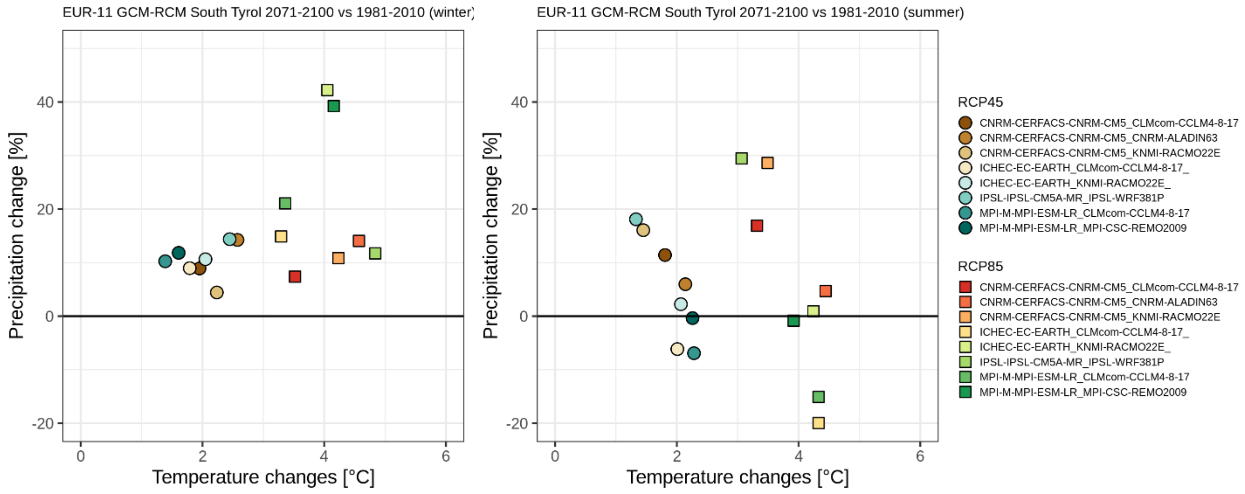

**Figure S1.** Mean temperature and precipitation changes in 2071-2100 from Euro-CORDEX models for South Tyrol under the RCP 4.5 (circles) and 8.5 (squares) scenario. Changes are computed with respect to the historical period 1981-2010.

Projections for solar radiation and relative humidity were available at a spatial resolution of 12 km, while temperature and precipitation were available at 1 km resolution. Therefore, all variables were post-processed in order to reach the resolution required by iLand. Solar radiation and relative humidity for these models were downloaded from the ESGF nodes (Cinquini et al., 2014) and the Copernicus Climate Data Store (Brönnimann et al., 2018). The EURO-CORDEX projections of solar radiation and relative humidity are split into several netCDF files and cover the entire European domain over a rotated-pole grid. We cropped and regridded them through bilinear interpolation onto the target 250-m grid covering the study region and we merged individual files into one single netCDF. These steps were performed by using the Climate Data Operator software (Schulzweida, 2023). For both variables, the ratio of the 1991-2020 monthly averages from resampled EURO-CORDEX and observations (downscaled EUMETSAT for solar radiation and downscaled EOBS for relative humidity) was computed for each grid point and used to bias-adjust the projections. All daily values from projections in the period 2006-2100 were multiplied for the correcting factor for the specific grid point and month. For relative humidity an additional control was added in order to assure that the correction did not lead to inconsistencies (RH > 100%). After the correction, all cases with RH > 100% were detected and replaced by 100. The same correction procedure was applied for temperature and precipitation. More specifically, the 1-km fields were resampled (bilinear interpolation) onto the higher resolution grid (250-m) and compared with the observations over the period 1991-2020. The monthly correction factors were derived and applied in order to adjust the EURO-CORDEX fields onto the finer orographic details of the target grid. The correction factors for minimum and maximum temperatures are additive and defined as the difference from observations. For each emission scenario (RCP45 and RCP85) we selected three climate models to cover the entire expected range of precipitation and temperature change (Figure S1), and we renamed the climate change projections from CC1 (most moderate) to CC6 (most extreme) scenario (Table S1, Figure S2).

**Table S1**. Name of the Euro-CORDEX models selected for the study (combination of global and regional model according to the institution that developed the simulations) and the corresponding name of the scenario used in this simulation experiment.

| **Euro-CORDEX model** | **Climate scenario** |
|---|---|
| CNRM-CERFACS-CM5_KNMI-RACMO22E | CC1 |
| CNRM-CERFACS-CNRM-CM5_CNRM-ALADIN63 | CC2 |
| MPI-M-MPI-ESM-LR_rcp45_r1i1p1_CLMcom-CCLM4-8-17 | CC3 |
| IPSL-IPSL-CM5A-MR_IPSL-WRF381P | CC4 |
| ICHEC-EC-EARTH_KNMI-RACMO22E | CC5 |
| ICHEC-EC-EARTH_CLMcom-CCLM4-8-17 | CC6 |

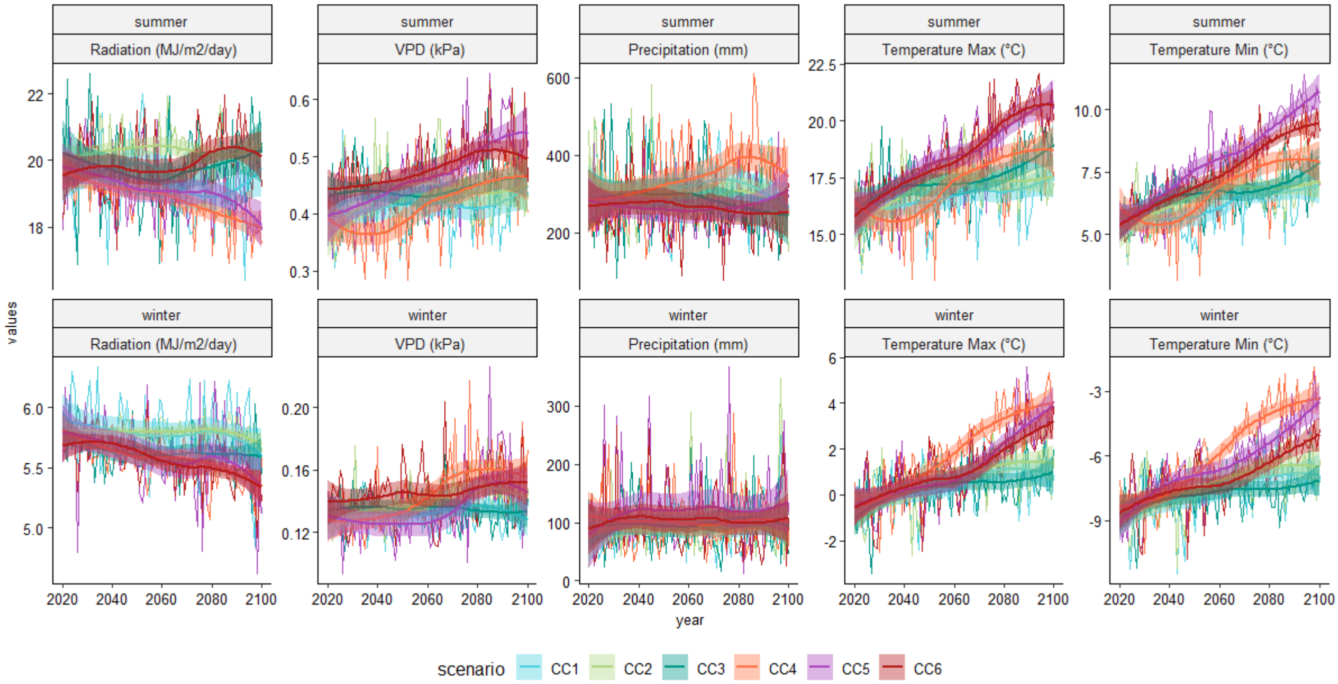

**Figure S2**. Trends of climate variables summarized over the study landscape for winter and summer seasons for the climate changes scenarios.

### PNV tree species

To generate a potential natural vegetation (PNV) forest landscape and to simulate its future dynamics, we restricted the species pool available in iLand to those species whose natural distribution ranges overlap with the study area. Following Caudullo et al., (2017) from the EUFORGEN project, we overlaid the study area with the species distribution maps provided in that work, resulting in a pool of 19 species (Table S2).

**Table S2**. List of the tree species used in the simulation study (left column) and those used for simulating PNV conditions relying on the natural distribution of tree species in our study area from the EUFORGEN project (right column).

| Available species in the study area (and in CF) | Presence in PNV simulation (PF) |
|---|---|
| *Abies alba* Mill. | Yes |
| *Acer campestre* L. | No |
| *Acer platanoides* L. | No |
| *Acer pseudoplatanus* L. | No |
| *Alnus glutinosa* (L.) Gaertn. | Yes |
| *Alnus incana* L. | Yes |
| *Alnus alnobetula* (Ehrh.) K.Koch | Yes |
| *Betula pendula* Roth, 1788 | Yes |
| *Betula pubescens* Ehrh., 1789 | Yes |
| *Carpinus betulus* L. | No |
| *Castanea sativa* Mill., 1768 | No |
| *Corylus avellana* L., 1753 | Yes |
| *Fraxinus excelsior* L., 1753 | Yes |
| *Larix decidua* Mill., 1768 | Yes |
| *Picea abies* (L.) H. Karst. | Yes |
| *Pinus cembra* L. | Yes |
| *Pinus mugo* Turra, 1764 | Yes |
| *Pinus nigra* J.F. Arnold | No |
| *Pinus sylvestris* L. | Yes |
| *Populus nigra* L. | No |
| *Populus tremula* L. | Yes |
| *Quercus petraea* (Matt.) Liebl. | No |
| *Quercus pubescens* Willd., 1805 | Yes |
| *Quercus robur* L., 1753 | No |
| *Salix caprea* L., 1753 | Yes |
| *Sorbus aria* (L.) Crantz | No |
| *Sorbus aucuparia* L. | Yes |
| *Tilia cordata* Mill. | Yes |
| *Tilia platyphyllos* Scop. | No |
| *Ulmus glabra* Huds. | Yes |

### Supplementary figures and tables

In this section, we reported the figures for the results of climate change scenarios not shown in the main manuscript (i.e. CC2, CC3, CC4, and CC5). First, for initial forest conditions (CF or PF) we calculated the coefficient of variation (CV) among scenarios replicates (10) with a 30-year time step starting from 2050 (see Table S2 below). CV was calculated on the basal area for each species within each stand cell (100 m resolution). Overall, we found low values of CV, indicating no important changes among simulation replicates.

**Tabel S3**. Coefficient of variation (in percentage) at a 30-year time step calculated for the 10 simulation replicates of every climate scenario using basal area output at the landscape scale for both PF and CF initialization conditions.

| Year | Initialization | baseline | CC1 | CC2 | CC3 | CC4 | CC5 | CC6 |
|---|---|---|---|---|---|---|---|---|
| 2050 | PF | 3.1 | 3.8 | 3.5 | 3.1 | 2.8 | 3.2 | 3.3 |
| 2080 | PF | 3.2 | 3.3 | 3.3 | 3.5 | 3.3 | 3.3 | 3.4 |
| 2110 | PF | 3.5 | 3.2 | 3.4 | 3.7 | 3.4 | 3.1 | 3.1 |
| 2140 | PF | 3.4 | 3.2 | 3.5 | 3.4 | 3.8 | 3.4 | 3.1 |
| 2170 | PF | 3.1 | 3.3 | 3.5 | 3.4 | 4.2 | 3.3 | 3.4 |
| 2200 | PF | 3.0 | 4.0 | 3.6 | 3.1 | 4.2 | 3.2 | 3.2 |
| 2050 | CF | 2.7 | 3.2 | 2.9 | 2.4 | 2.6 | 2.9 | 2.8 |
| 2080 | CF | 3.1 | 4.1 | 3.8 | 3.5 | 4.1 | 3.8 | 3.4 |
| 2110 | CF | 3.3 | 4.4 | 4.2 | 4.2 | 4.3 | 4.5 | 4.0 |
| 2140 | CF | 3.4 | 4.4 | 4.5 | 3.6 | 4.7 | 4.8 | 3.0 |
| 2170 | CF | 3.2 | 4.1 | 4.2 | 3.5 | 4.3 | 5.2 | 2.6 |
| 2200 | CF | 3.4 | 3.9 | 3.8 | 3.7 | 4.9 | 5.4 | 2.9 |

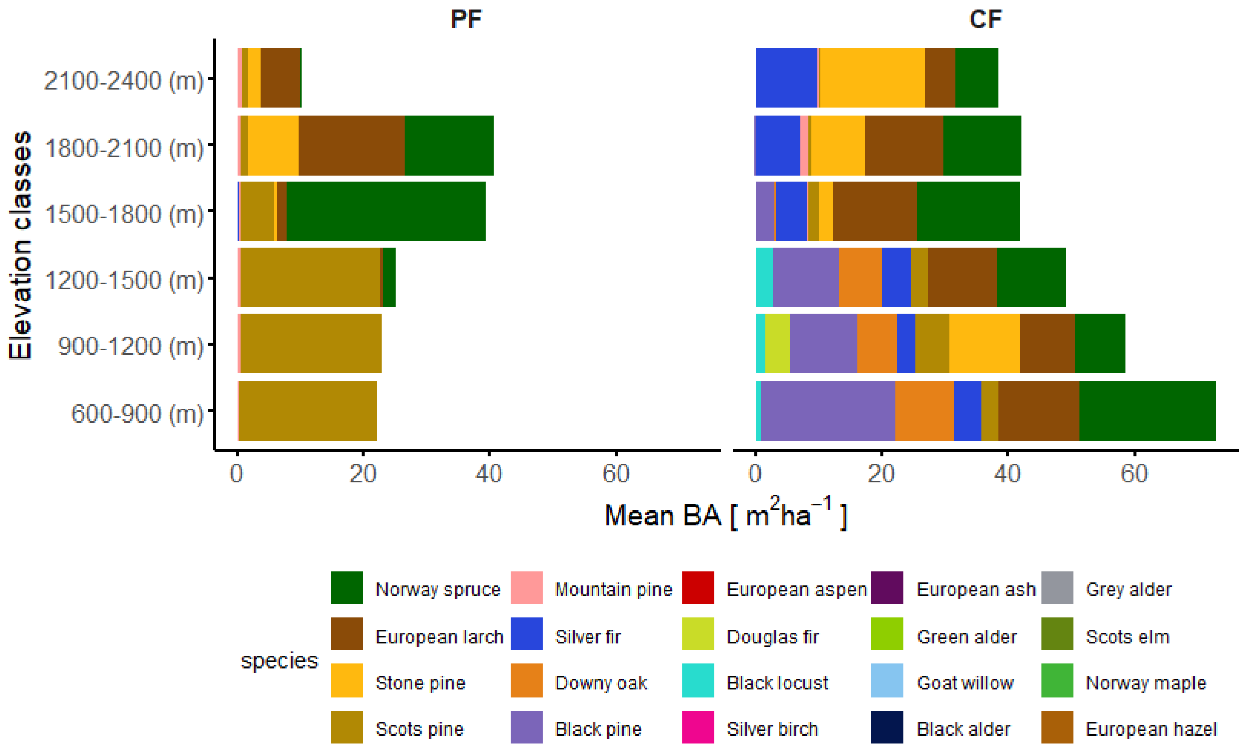

**Figure S3.** Mean species share in terms of basal area across elevation classes for the two initialized landscapes PF and CF. It is important to note that at lower elevation (< 1200 m asl), PF presents a higher diversity which is not observable due to the low share of the minor species. On the contrary, in CF those elevations appear more diverse due to the larger share of the different species.

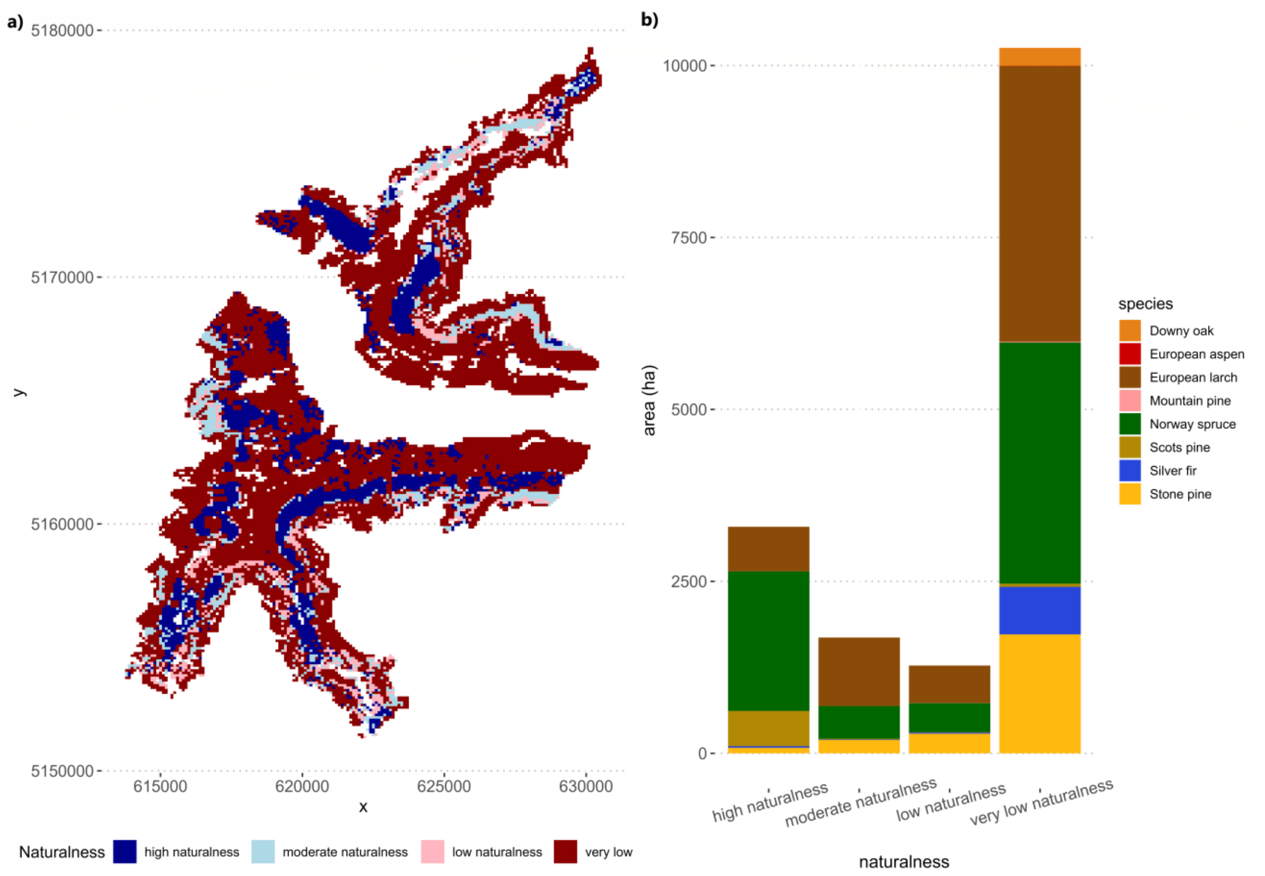

**Figure S4.** Panel (a): the map of the naturalness score divided into 4 categorical classes equally divided by a 0.25 score-step: 1) high naturalness, for $NAT_{score}$ = 0.75 - 1; 2) moderate naturalness, for $NAT_{score}$ = 0.50 - 0.70; 3) low naturalness, for $NAT_{score}$ = 0.25 - 0.50; 4) very low naturalness, for $NAT_{score}$ < 0.25, going from red (very low naturalness) to blue (high naturalness). Panel (b): extension of each naturalness category over the landscape and its characterization in terms of dominant species.

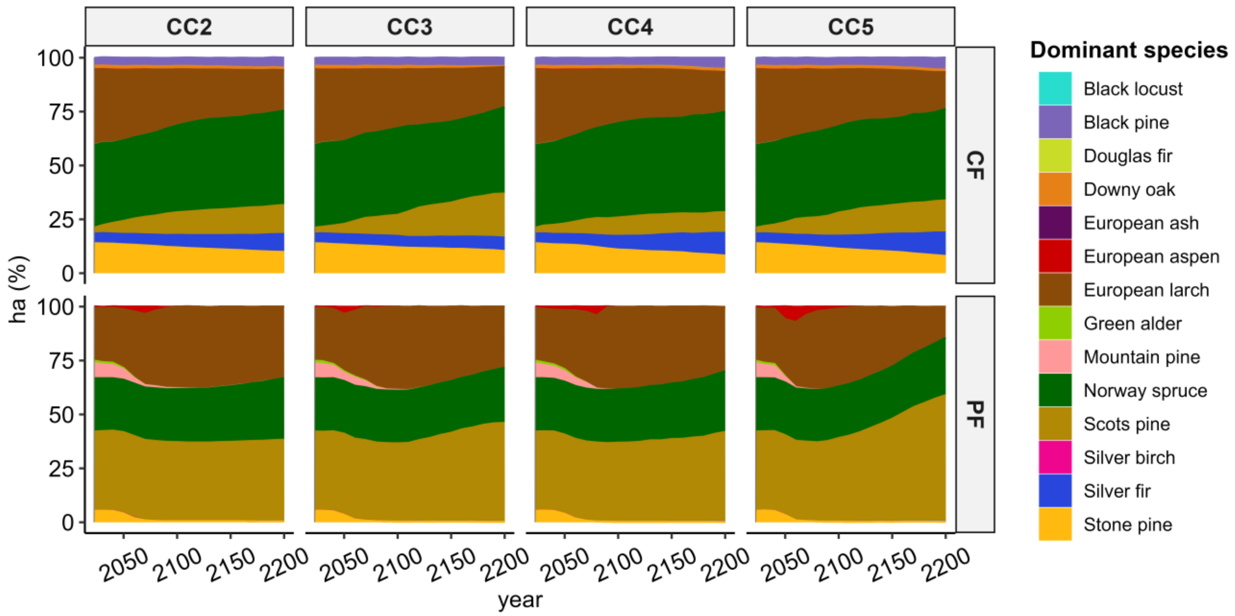

**Figure S5**. Dominant species share at the landscape level (percentage of ha) for CF and PF under future projections of historic CC2, CC3 (RCP 4.5), CC4, and CC5 (RCP 8.5) until 2200.

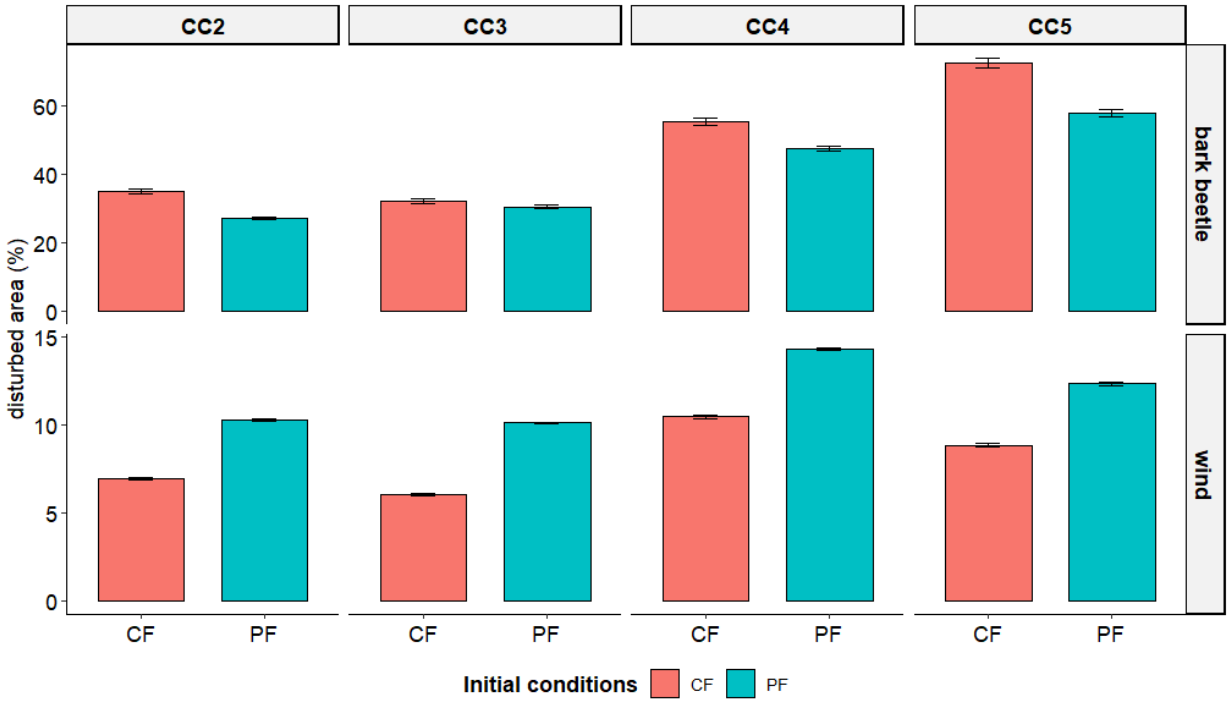

**Figure S6**. Impact of bark beetle (upper panel) and wind (lower panel) on PF (red) and CF (blue) under CC2, CC3, CC4, and CC5. Bars represent the cumulative impact across the entire simulation period. Error bars show the standard deviation of the replicates mean. Note the different scales of the y axis for the two disturbance agents. Bark beetle and wind damages were analyzed using the outputs of the corresponding disturbance modules in iLand, which provide several spatial variables. For bark beetles, we summed the variables infestedArea_ha and killedArea_ha. The former represents all newly infested cells, while the latter includes only those cells in which bark beetle activity resulted in tree mortality. For wind disturbance, we used the only available spatial variable, area_ha, which similarly represents the cells where wind caused tree mortality. Further details on the disturbance outputs are available in the iLand web-based documentation for the wind (https://iland-model.org/wind+module) and bark beetle modules (https://iland-model.org/barkbeetle+module). For both disturbance types, outputs were recorded annually at the landscape scale. It is important to note that disturbance impacts are simulated at the stand level; therefore, the total damaged area may be slightly overestimated if only a portion of a pixel was actually affected.

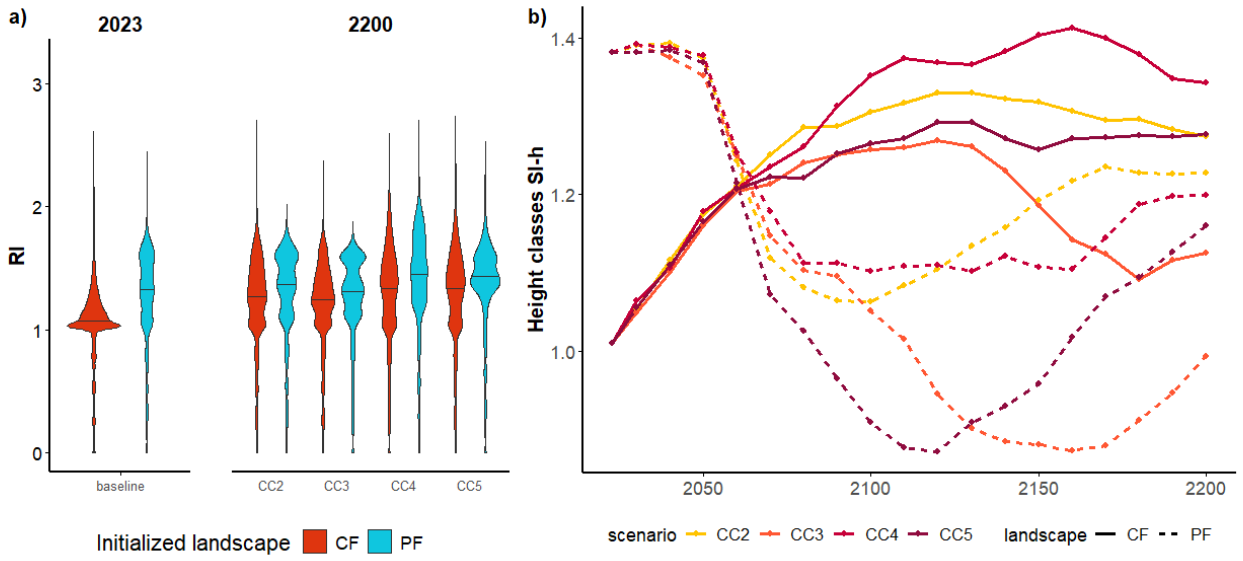

**Figure S7**. Panel (a) Violin plots showing the rumple index (values of the distribution across the forested landscape extracted at 1 ha resolution) comparing PF and CF at the beginning and the end of the simulation under baseline, CC2, CC3, CC4, and CC5 scenarios (median: horizontal black line). Panel (b): Temporal trends of the Shannon Index of the height classes, representing the evenness of the stand height classes distribution plotted over time with time steps of 10 years. Higher values indicate greater diversity in vertical stand structure.
